# Developmental NMDA receptor signaling regulates cerebellar unipolar brush cell number and dampens excitability

**DOI:** 10.64898/2026.08.21.744536

**Authors:** Harsh N. Hariani, Gabrielle G. Peña, Zoey E. Joshlin, Timothy S. Balmer

**Affiliations:** School of Life Sciences, Arizona State University, Tempe, AZ, USA

**Author notes:** **Corresponding Author:** Timothy Balmer.

## Abstract

Unipolar brush cells (UBCs) are excitatory interneurons that have a characteristic dendritic brush that amplifies and extends incoming signals in the cerebellum. UBCs transform synaptic input through their ionotropic and metabotropic glutamate receptors. Differential regulation of receptor subunits is a critical developmental process, but how the expression of glutamatergic receptors changes in UBCs as they develop is unclear. NMDA-type glutamate receptors (NMDARs) are particularly important for development and plasticity. We examined the expression of NMDAR subunits during development and tested whether signaling through these receptors is necessary for the development of the elaborate dendritic structure and unusual synaptic function of UBCs. Whole-cell patch clamp recordings from UBCs in acute brain slices revealed tonic and synaptic NMDAR-mediated currents in early postnatal UBCs that decrease during development. RNAscope in situ hybridization revealed differential developmental regulation of GluN2C/D subunits. Cell-type specific constitutive NMDAR knockout had no apparent effect on dendritic brush development, but increased UBC number in adulthood, suggesting a role in programmed cell death. Both pharmacological blockade or genetic deletion of NMDARs produced a paradoxical increase in excitability, which was calcium dependent and was occluded by inhibition of calcium activated potassium channels. Thus, NMDA receptors are dispensable for migration and dendritic development but may be involved in cell death pathways. Their functional roles include synaptic signaling as well as providing a tonic calcium flux that dampens excitability in developing UBCs and may influence transformations of vestibular signals essential for smooth movements and balance.

**SIGNIFICANCE STATEMENT:** Unipolar brush cells (UBCs) amplify and transform incoming sensorimotor signals to support cerebellar function. Their disruption could lead to disorders such as ataxia and nystagmus. Factors contributing to UBC development are unclear. NMDA receptors play a pivotal role in maturation of other cerebellar neuron types. Here we show that NMDA receptors are expressed by developing UBCs and that they regulate the number of UBCs that survive through development, but do not contribute to their dendritic brush morphology. These receptors generate a tonic current due to their GluN2C/D subunit expression that paradoxically dampens excitability through the activation of calcium activated potassium channels. Thus, the expression of NMDA receptors in these neurons is a mechanism to regulate their excitability and their number.

## INTRODUCTION

N-methyl D-Aspartate (NMDA) receptors (NMDARs) are present throughout the central nervous system across species and play important roles in cellular and circuit development (Forrest et al., 1994; Kalb, 1994; Rajan and Cline, 1998; Lee et al., 2005; Namba et al., 2011; Gao et al., 2018; Hanson et al., 2019; Hansen et al., 2021). The vestibulocerebellum is essential for maintaining body posture, head position, and eye reflexes (Miles, 1987; Manzoni, 2007; Prati et al., 2024). In the vestibulocerebellar cortex, the two glutamatergic interneurons, granule cells and unipolar brush cells (UBCs), express NMDARs (Rossi et al., 1995; Cathala et al., 2000; Billups et al., 2002). Granule cells require NMDARs for migration, cell survival and dendritic development and circuit maturation (Komuro and Rakic, 1993; Hirai and Launey, 2000).

In contrast, the role of NMDARs in UBCs remains unclear. UBCs are predominantly located in the vestibular cerebellum and are present across vertebrates, including humans (Rossi et al., 1995; Diño et al., 1999; Takács et al., 1999; Víg et al., 2005). They receive input from numerous sources including the vestibular ganglion and from the medial vestibular nucleus (Balmer and Trussell, 2019). UBCs vary in their response to synaptic stimulation, ranging from ON (increase in firing) to OFF (pause in firing) (Borges-Merjane and Trussell, 2015; Guo et al., 2021; Huson et al., 2023; Huson and Regehr, 2024). The structure of the UBC dendritic brush is suggested to contribute to their amplification of synaptic responses (Kinney et al., 1997; Mugnaini et al., 2011). Factors governing dendritic brush development are not known, despite their importance to synaptic function. Given the general importance of NMDARs in neuronal development, we hypothesized that NMDAR activity contributes to the development of UBCs.

NMDARs consist of tetraheteromers containing a combination of subunits that determines their glutamate affinity, kinetics, ion permeability, and downstream signaling pathways (Cull-Candy and Leszkiewicz, 2004; Vyklicky et al., 2014; Haddow et al., 2022). Differential regulation of these subunits through development is necessary for normal cell morphology and synaptic development (Akazawa et al., 1994; Monyer et al., 1994; Sheng et al., 1994; Cathala et al., 2000; Henson et al., 2008). Typically, activation of NMDARs requires binding of agonists with coincident depolarization, to relieve the magnesium pore block. However, NMDARs with GluN2C/D subunits can mediate currents at hyperpolarized potentials owing to their relative insensitivity to magnesium (Hollmann and Heinemann, 1994; Qian et al., 2005; Paoletti et al., 2013). Moreover, GluN2C/D subunits can mediate a tonic current which is necessary for promoting dendritic and synaptic maturation, and regulation of neuronal excitability (Gall and Dupont, 2019; Hanson et al., 2019). UBCs are exposed to ambient glutamate that produces tonic AMPA and mGluR2 currents (Balmer et al., 2021), but it is not known whether they have a tonic NMDAR-mediated current.

In granule cells, NMDAR subunit composition is developmentally regulated from GluN2B to GluN2A and eventually GluN2C in adulthood, presumably to meet the needs of the developing granule cell (Cathala et al., 2000). It is not known if UBCs express NMDARs at all ages and whether they are expressed in some UBC subtypes (Canton-Josh et al., 2022). Most studies using mice over P21 either do not address NMDAR-mediated currents or report that NMDARs had no significant contributions to the synaptic response at the MF– UBC synapse (Borges-Merjane and Trussell, 2015; Hariani et al., 2024; Huson and Regehr, 2024). While NMDARs appear to be developmentally regulated in UBCs, their role in development and synaptic function is unclear.

Here, we used a subset of ON UBCs as a model to study the developmental role and physiological function of NMDARs in UBCs. We confirmed the presence of functional NMDARs and determined their subunit composition across development using whole-cell patch clamp electrophysiology and *in situ* hybridization. Deleting NMDARs from UBCs revealed that they regulate UBC number, but they are not necessary for dendritic brush development. Tonically conducting GluN2C/D containing NMDARs dampen excitability through their activation of calcium activated potassium channels.

## METHODS

### Animals

We used transgenic mice of both sexes. The GRP line (Tg(Grp-Cre)KH107Gsat, MMRRC_031182-UCD) expresses cre recombinase under the GRP (gastrin releasing peptide) promoter (Gerfen et al., 2013). The Ai9 line (Gt(ROSA)26Sor^tm9(CAG-tdTomato)Hze^, MSR_JAX:007909) expresses tdTomato in a cre-dependent manner (Madisen et al., 2010). A cross between these two lines (GRP/Ai9) has previously been shown to label a subset of mGluR1(+) ON UBCs in the cerebellum. The P079 line Et(tTA/mCitrine)P079Sbn has been previously shown to label most calretinin(+) OFF UBCs in the cerebellum with mCitrine (Shima et al., 2016; Hariani et al., 2024). GluN1 knock-out (KO) animals were purchased from Jackson Laboratories (#005246; B6.129S4-GluN1^tm2Stl/J^) and bred with our GRP and Ai9 mice to generate GluN1^flox/flox^/GRP^+/−^/Ai9^+/−^ mice (Tsien et al., 1996). Transgenic mice were genotyped by PCR. For PPDA experiments, wild-type C57BL/6J mice of age P10-14 were used. For synaptic stimulation experiments, C57BL/6J-TgN(grm2-IL2RA/GFP)1kyo (mGluR2-GFP) line of age P10-14 were used (Nunzi et al., 2002; Borges-Merjane and Trussell, 2015; Balmer and Trussell, 2019). To confirm that they were ON UBCs, we confirmed an increase in firing following either glutamate puff application or synaptic stimulation of MFs in current clamp configuration. From **Fig** 2 onward, all experiments were conducted on mGluR1(+) ON UBCs labeled in the GRP line.

For developmental comparisons, P10-14 or P25-30 mice were used. UBCs reach the internal granular layer by ∼P8 and their brush is still immature and developing (Morin et al., 2001; Englund et al., 2006). P15-P16 is the critical period for climbing fiber development (Kakizawa et al., 2000). Granule cells finish migrating into the granular layer by P16 and cerebellar foliation patterns also mature by P16 (Leto et al., 2016). UBC brush morphology and receptor physiology reaches maturity at P21-28 and cerebellar lobules reach synaptic and anatomic maturity at ∼P20 (Morin et al., 2001; White and Sillitoe, 2013). For these reasons, we chose P10-14 as our ‘developing’ and P25-30 as our ‘mature’ time points.

### Acute brain slice preparation

Mice were anaesthetized using isoflurane and the brain was extracted in ice-cold sucrose-artificial cerebrospinal fluid (ACSF) containing (in mM): 87 NaCl, 75 sucrose, 25 NaHCO_3_, 25 glucose, 2.5 KCl, 1.25 NaH_2_PO_4_, 0.4 Na-ascorbate, 2 Na-pyruvate, 0.5 CaCl_2_, 7 MgCl_2_, bubbled with 5% CO_2_/95% O_2_. 250 - 300 µm sagittal cerebellar slices were prepared with a vibratome (Campden Instruments). Slices were incubated in recording ACSF at 35°C for 30 minutes and then stored at room temperature. Recording ACSF solution contained (in mM): 130 NaCl, 2.1 KCl, 1.2 KH_2_PO_4_, 3 Na-HEPES, 10 glucose, 20 NaHCO_3_, 0.4 Na-ascorbate, 2 Na-pyruvate, 1.5-2.4 CaCl_2_, 1 MgSO_4_, bubbled with 5% CO_2_ / 95% O_2_ (300–305 mOsm). Slices were used for electrophysiology within 8 hours of slicing. Recordings were made from lobe X of the cerebellum.

### Electrophysiology

Brain slices were perfused with ACSF at 2-3 ml/min using a peristaltic pump (Ismatec). The ACSF bath was maintained at 32-34°C using an inline heater (Warner Instruments). Synaptic inhibition was blocked by 0.5 µM Strychnine and 5 µM SR95531. PPDA experiments were conducted at room temperature. A horizontal pipette puller (Sutter Instruments) was used to make borosilicate glass pipettes with open tip resistances of 6-8 MΩ. Potassium-based internal solution contained (mM): 113 K-gluconate, 9 HEPES, 4.5 MgCl2, 0.1 EGTA, 14 Tris-phosphocreatine, 4 Na2-ATP, 0.3 tris-GTP, 0.1-0.3% biocytin, supplemented with either 1 µM Alexa Fluor 488 or 594. Cesium-based internal solution contained (in mM): 145 Cs-methylsulfonate, 10 QX-314, 2 MgCl2, 5 K2ATP, 0.5 EGTA, 5 HEPES. Osmolality was adjusted to 290 mOsm using sucrose. pH was adjusted to 7.2-7.25 using KOH or CsOH for potassium or cesium based internal solutions, respectively. Cells were voltage clamped at −75 mV. 15 mV (K-gluconate) and 12 mV (Cs-methylsulfonate) junction potential correction was applied to all reported voltages. Data were acquired with a Multiclamp 700B amplifier and pClamp 11 software and quantified using Clampfit software (Molecular Devices). Signals were sampled at 50-100 kHz using a Digidata (1550B, Molecular Devices) digitizer. Electrical stimulation was performed by white matter stimulation of 1-90 V (10 stimulation train at 50 Hz) using a concentric bipolar microelectrode (FHC; 30202). Glutamate puff experiments were performed using 1 mM monosodium glutamate solution in ACSF in a pipette and 20-30 ms duration applications at 5-10 PSI. For current clamp recordings, bias current was adjusted to maintain the cell at similar membrane potentials before and after drug application. For all cells, membrane potential was maintained between −77 mV to −85 mV. For each cell, trials only within a tight range of 3.7 ± 0.42 mV were used for analysis. Trials that deviated more than that before and after drug application were excluded. Latency to first spike was measured from the start of first stimulation to the abrupt increase in the rate of depolarization. Duration of burst was measured from the abrupt increase in the rate of depolarization to the abrupt decrease in the rate of hyperpolarization. Input resistance was measured using Clampfit software and by the slope of I-V curves measured when the membrane potential reached steady state during hyperpolarizing current steps.

### RNAscope

RNAscope was performed using the RNAscope Multiplex Fluorescent V2 Assay (Advanced Cell Diagnostics, catalog # 323280). P7, P12 and P30 aged mice from the GRP/Ai9 mouse line were used. Brains were extracted in ice-cold sucrose ACSF and drop-fixed in 4% PFA overnight at 4°C. Brains were then sequentially incubated in 10%, 20% and 30% sucrose solution in PBS at 4°C, changing from one solution to the next once the brain sank to the bottom of the container. Brains were flash-frozen with ethanol on dry ice in optimal cutting solution (OCT) (Tissue-Tek; 62550-01) and stored at −80°C. Brains were sliced sagittally at 15 µm using a cryostat and mounted onto superfrost plus slides (Fisherbrand; 12-550-15) and stored at −80°C until further processing. Brains from 3-4 mice per condition were used. For each brain, five representative sections from medial lobe X of the cerebellum were mounted. Lab surfaces were cleaned with RNaseZAP (Sigma Aldrich; R2020) before running the assay. RNAscope assay was performed following the manufacturer’s instructions and recent work that used the same probes (Drotos et al., 2025). Briefly, slides were washed with PBS and then baked for 30 minutes at 60°C followed by post-fixation in 4% PFA in PBS for 15 minutes at 4°C. Tissue on slides were dehydrated with sequential washes in increasing concentrations (50% - 100%) of ethanol. Slides were then incubated in hydrogen peroxide for 10 minutes at room temperature followed by performing antigen retrieval as per manufacturer’s guidelines. Slides were dried with a Kim wipe and hydrophobic barriers were drawn around sections. Slides were incubated in RNAscope Protease III for 30 minutes at 40°C. Control slices were incubated with the positive and negative probes (1 slice each) (Catalog# PN 320881 and PN320871, respectively) while all other slices were incubated with tdTomato, GluN2C and GluN2D probes (Catalog# 317041, 445581-C2 and 425951-C3, respectively) for 2 hours at 40°C. Next, probe amplification was performed by separate incubation with each AMP (1,2 and 3) (included in kit) for 30 minutes (AMP1, 2) or 15 minutes (AMP3) at 40°C. Signal was developed by incubation with HRPC1-3 (included in kit) for 15 minutes at 40°C and 3 different TSA vivid fluorophores were used for labeling probes C1-3 by incubation with diluted fluorophores for 30 minutes at 40°C (tdTomato was labeled with TSA Vivid 570 (323272), GluN2C with TSA Vivid 520 (323271) and GluN2D with TSA Vivid 650 (323273)). Slides were then counterstained with DAPI (included in kit) by incubation for 30 seconds at room temperature followed by coverslipping (Fisherbrand; 12544D) using ProLong Gold Antifade mountant (Invitrogen; P36930). Slides were cured overnight at room temperature and imaged the next day.

Slides were imaged with Olympus VS200 slide scanner using a 40X objective. Acquisition settings were identical for all slides. Exposure times were set to 50, 100, 70, and 50 ms for DAPI, FITC, TRITC and Cy5, respectively. Images were analyzed using QuPath software (Bankhead et al., 2017). The tdTomato signal was used to segment UBC cell bodies by running the automated segmentation pipeline using the ‘cell detection’ feature. Next, GluN2C and GluN2D puncta were segmented using the ‘subcellular detection’ feature. Segmentation was manually verified to ensure specificity and sensitivity. Thresholding values were conservatively set based on pixel intensities of the three channels from the negative control samples. Default spot and cluster parameters were used. ‘Estimated spots/cell’ and ‘mean fluorescence intensity/cell’ were used as primary readouts. These readouts quantified the number of puncta colocalized with the tdTomato signal and the mean fluorescence of intensity within the segmented tdTomato cell body, respectively, for each channel representing GluN2C and GluN2D.

### UBC number and UBC brush analysis

Mice from 2 experimental groups were used - GRP/Ai9 control mice and GluN1 KO GRP/Ai9 mice. Brains were extracted from 3 animals of each group aged P12 into ice-cold sucrose ACSF and drop-fixed in 4% PFA overnight at 4°C. 3 animals from each group of age P30 were perfused with 4% PFA and further post-fixed in 4% PFA overnight at 4°C. Cerebella from all brains were then sectioned sagittally at 50 µm thickness using a vibratome (Campden Instruments). Every other slice was mounted on superfrost plus slides (Fisherbrand; 12-550-15) along the mediolateral axis. Slides were coverslipped (Fisherbrand; 12544D) with Fluoromount-G (Southern Biotech) which included DAPI as a counterstain. Slides were cured overnight at room temperature and imaged the next day with Olympus VS200 slide scanner using a 10X objective. Image analysis was performed using the QuPath software (Bankhead et al., 2017). A region of interest was drawn around the granular layer of lobe X using the ‘polygon annotation’ tool and labeled UBCs were counted manually using the ‘add points to annotation’ feature. The number of UBCs per slice and density of UBCs/mm^2^ were used as readouts. To count UBCs in the dorsal cochlear nucleus, 5 equally spaced coronal slices along the rostrocaudal axis from the dorsal cochlear nucleus on one side were used for analysis. UBC counting was performed the same way as for the cerebellum.

For UBC brush volumetric analysis, individual UBCs were imaged using a confocal microscope (LSM800, Zeiss) with Airyscan processing which performs deconvolution post-acquisition to increase signal-to-noise ratio. UBCs imaged were sampled from at least 3 animals and from at least 5 slices/animal for each experimental group. Images were analyzed using Arivis4D (Zeiss). We performed intensity-based segmentation and rendered a 3D reconstruction for each cell. Each 3D cell rendering was sliced into two distinct brush and soma volumes. Brush volume and surface area were used as readouts.

### Computational model

A single compartment model was built using NEURON (Hines and Carnevale, 1997; Carnevale, 2005) and included voltage-gated sodium (gNa) and potassium (gK) conductances to produce action potentials (Destexhe et al., 1994), a passive leak conductance (gpas), a calcium activated potassium conductance (gBK) and an NMDA receptor conductance (gNMDA), described below. The model cell was 20 μm in diameter and had the specific membrane capacitance of 1 μF/cm^2^. Temperature for all mechanisms was set to 37°C.

|  |  |
| --- | --- |
| gNa (S/cm<sup>2</sup>) | 0.01 |
| gK (S/cm<sup>2</sup>) | 0.008 |
| gBK (S/cm<sup>2</sup>) | 0.01 |
| gNMDA (S/cm<sup>2</sup>) | 0.001 |
| gpas (S/cm<sup>2</sup>) | 0.0001 |
| ENa (mV) | 50 |
| EK (mV) | -77 |
| Epas (mV) | -70 |

### Tonic NMDA receptor conductance

The NMDA current was treated as a non-specific cation current with 10% of the current carried by calcium (Schneggenburger et al., 1993; Burnashev et al., 1995) using the following equation:

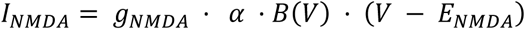

Where *g_NMDA_*is the maximal NMDA conductance density, *α* is a scaling factor representing the level of tonic receptor activation from 0 to 1, *E_NMDA_*is the reversal potential (0 mV), and *B(V)* is the voltage-dependent magnesium block and was implemented using the formulation of (Jahr and Stevens, 1990) fit to the data shown in **Fig** 1C (Developing (P10-14) ON UBCs):

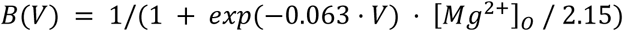

**Figure 1:**
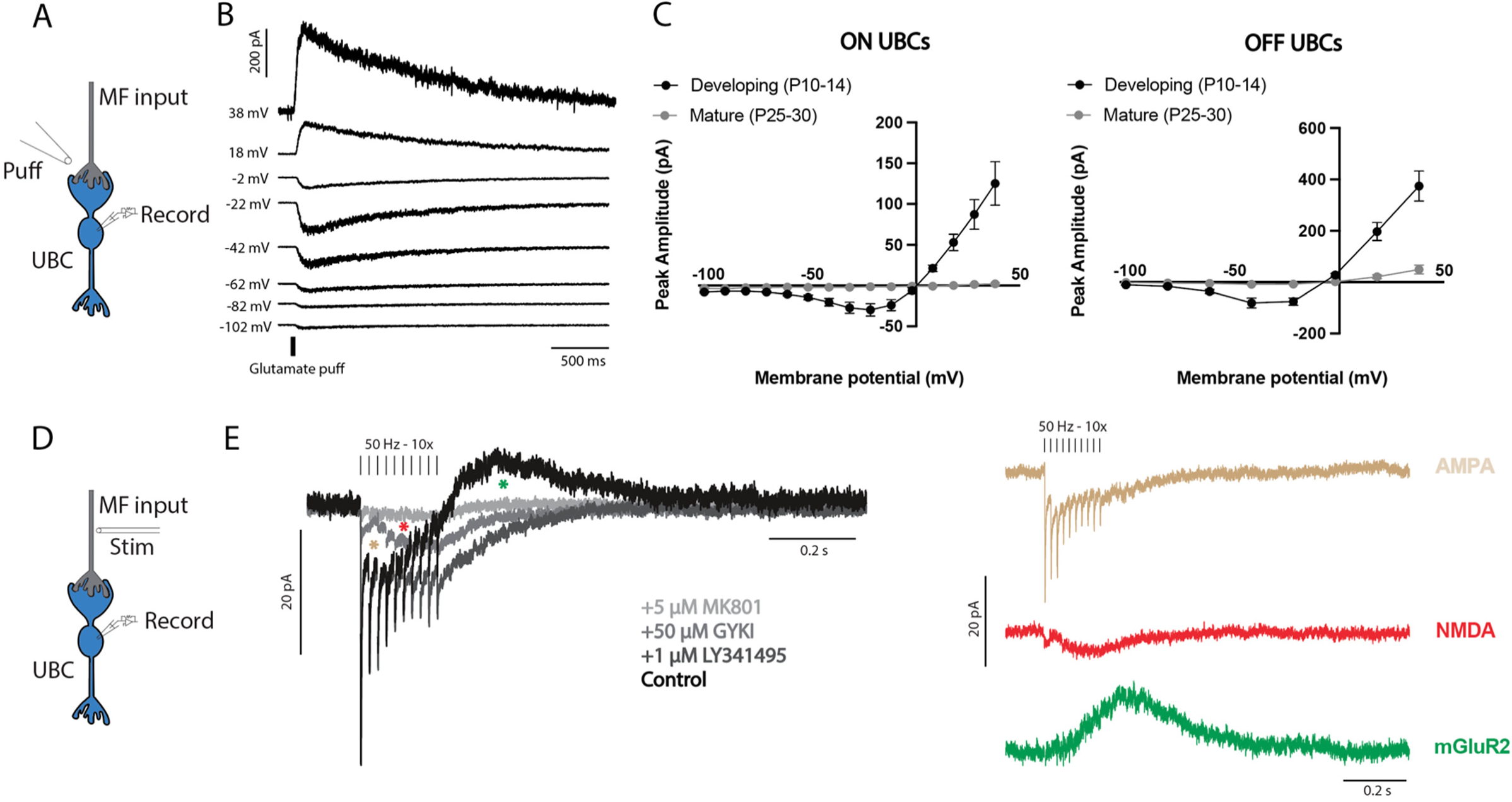
NMDAR-mediated currents in developing cerebellar UBCs A) Illustration of experimental design showing brief (10-30 ms) glutamate puff on UBC brush while recording from soma. B) NMDAR-mediated currents increased on depolarization before reversing near the expected 0 mV. Note the presence of NMDAR-mediated currents even at hyperpolarized potentials. Traces are from a developing P12 ON UBC. C) NMDA I-V curves from glutamate puff evoked currents show a range of negative slope conductance in ON (left) and OFF (right) UBCs. Developing (P10-14) UBCs had large NMDAR-mediated currents that were nearly absent in mature (P25-30) ON UBCs and greatly reduced in mature OFF UBCs. D) Illustration of experimental design showing MF stimulation while recording from UBC soma. E) Synaptic currents elicited by MF stimulation. Subtractions of currents before and after applying antagonists enabled isolation of individual receptor contributions to the total evoked current (AMPA- beige, NMDA- red and mGluR2- green), confirmed the presence of synaptic NMDA receptors.

The calcium component contributed to the intracellular [Ca^2+^]_i_ used by the calcium accumulation equation and the BK conductance described below. Because the receptor is treated as tonically active, no explicit glutamate-binding kinetics or desensitization were included in order to capture the steady-state effect of ambient extrasynaptic NMDA receptor activation rather than synaptically evoked events.

### Intracellular calcium dynamics

Calcium accumulation was modeled using a single-shell approximation (Destexhe et al., 1994):

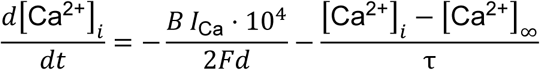

with shell depth *d* = 0.1 µm, removal time constant τ = 80 ms, resting [Ca^²*^]_i_ = 50 nM, buffering factor (*B*) = 1, and (*F*) Faraday’s constant. *I_Ca_* includes the calcium current through the NMDAR. The resulting [Ca²*]_i_ gates the BK conductance. τ = 80 ms is the time constant of calcium removal, and [Ca²*]_∞_ = 50 nM is the resting intracellular calcium concentration. The factor of 10⁴ converts current density (mA/cm²) and shell depth (µm) into consistent units of mM/ms, and the factor of 2 accounts for the divalent charge of calcium. The shell volume scales with the diameter (*d*) of the compartment.

### Calcium activated potassium conductance (BK)

A simplified Calcium activated potassium conductance was used to approximate BK conductance (Destexhe et al., 1994). Activation depended only on [Ca²*]_i_ using a Hill function with coefficient 2 and half-activation [Ca²*]_1/2_ = 0.1 µM and forward and backward rates of 0.1 ms^−1^. The variable *o* represents the fraction of BK channels in the open state.

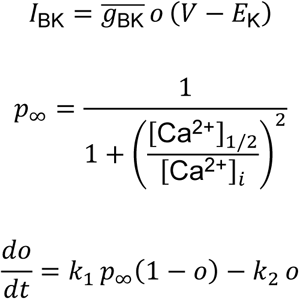

To generate **Fig** 7B-D, a parameter sweeps from 0.00005 to 0.0005 S/cm^2^ conductance values for gNMDA and gBK was performed and latency to first spike, rise slope of depolarization, and number of spikes were calculated and plotted in 21×21 grids.

## RESULTS

### NMDAR-mediated currents in developing cerebellar UBCs

To gain a broad understanding of NMDAR function in UBCs at different ages, we recorded from ON and OFF UBCs in developing and mature animals using whole-cell patch-clamp electrophysiology while stimulating the cell with brief focal application (puff) of 1 mM glutamate (**Fig** 1A). In the presence of AMPA, mGluR1 and mGluR2/3 glutamate receptor antagonists (50 μM GYKI53655, 1 μM JNJ16259685 and 1 μM LY341495, respectively), a glutamate puff produced robust NMDAR-mediated currents. The peak current amplitude increased with membrane depolarization from ∼ −80 mV to ∼ −20 mV, decreased above −20 mV, and then reversed at ∼0 mV (**Fig** 1B). Current – voltage (I-V) curves for developing ON (black, left, n = 12) and OFF (black, right, n = 7) are characteristic of NMDAR-mediated currents, having a negative slope conductance between −80 and −20 mV (**Fig** 1C-D) (Mayer et al., 1984; Sah et al., 1989; Chiu and Carter, 2022). We observed a near-complete lack of NMDAR-mediated currents in mature ON UBCs (grey, left, n = 12) while mature OFF UBCs (grey, right, n = 11) showed a profound reduction in NMDAR-mediated currents (at +38 mV: ON UBCs: Developing (125.3 ± 26.82 pA, n = 11) vs Mature (2.15 ± 1.65 pA, n = 8), Mann-Whitney test; p < 0.0001; OFF UBCs: Developing (374.3 ± 58.12 pA, n = 7) vs Mature (49.34 ± 16.84 pA, n = 8), Welch’s test; p = 0.0010; mean ± SEM).

Electrical stimulation of presynaptic MFs evoked synaptic currents in developing ON UBCs (**Fig** 1D, E). Sequential application of AMPA, mGluR2/3 and NMDA (10 μM MK801) receptor antagonists and subsequent subtraction allowed us to identify individual receptor contributions to the total current elicited (**Fig** 1E) (NMDAR-mediated current area: 1163.55 ± 356.18 fC, n = 6; mean ± SEM). These experiments demonstrated the presence of functional NMDARs in developing ON UBCs which can be activated by synaptic release of glutamate.

Thus, developing UBCs have functional NMDA receptors that produce large currents that decrease significantly during maturation. Since ON UBCs showed the most dramatic differential regulation of NMDARs across development, we decided to use them as a model to understand the role of NMDARs in developing UBCs in the following experiments.

### Developing ON UBCs express GluN2C/D subunits that produce a tonic current

We observed that UBCs had NMDAR-mediated currents at hyperpolarized potentials that were unlikely to be mediated by GluN2A/B due to their pore block by extracellular magnesium. We hypothesized that developing ON UBCs expressed GluN2C/D subunits which are relatively insensitive to the magnesium block and can therefore be active at hyperpolarized potentials. To explore the subunit composition of NMDARs in developing ON UBCs, we first used the well-characterized pharmacological agent, Ifenprodil, to test for the presence of GluN2A/B. Ifenprodil is a selective antagonist for GluN2B and heteromeric GluN2A/B receptors at 10 μM (Kew et al., 1996; Cathala et al., 2000). Using glutamate puffs we generated I-V curves before and after application of 10 μM Ifenprodil (**Fig** 2A, left). The black trace represents glutamate puff evoked NMDAR-mediated currents. Either Ifenprodil (blue trace) or MK801 (red trace) was applied, blocking the majority of the current (**Fig** 2A, left). Inhibition of the current by Ifenprodil was voltage dependent, as the proportion of residual (unblocked) current depended on membrane potential (**Fig** 2A, right). Ifenprodil only blocked a small proportion of the current at −102 mV, having a larger residual current of (74.88 ± 11.81 %; mean ± SEM) compared to MK801 (21.64 ± 10.28 %; mean ± SEM) (n = 6; Unpaired t-test; p = 0.0068). The residual current was significantly larger for Ifenprodil compared to MK801 at other potentials (n = 6; Unpaired t-test; mean ± SEM): - 62 mV (Ifenprodil: 36.46 ± 6.537 %; MK801: 14 ± 4.173 %; p = 0.0160), −22 mV (Ifenprodil: 17.80 ± 4.643 %; MK801: 4.563 ± 2.149 %; p = 0.0271) and at +38 mV (Ifenprodil: 9.756 ± 3.260 %; MK801: 1.973 ± 0.5144 %; p = 0.0289). The effect varied significantly across membrane potentials for Ifenprodil (n = 6; One-way ANOVA; p < 0.0001), but not for MK801 (n = 6; One-way ANOVA; p = 0.0849). These results suggest that although GluN2A/B subunits conduct the majority of the current at depolarized potentials, the presence of Ifenprodil insensitive currents at hyperpolarized potentials supports the hypothesis that the current is partly mediated by GluN2C/D subunits.

**Figure 2:**
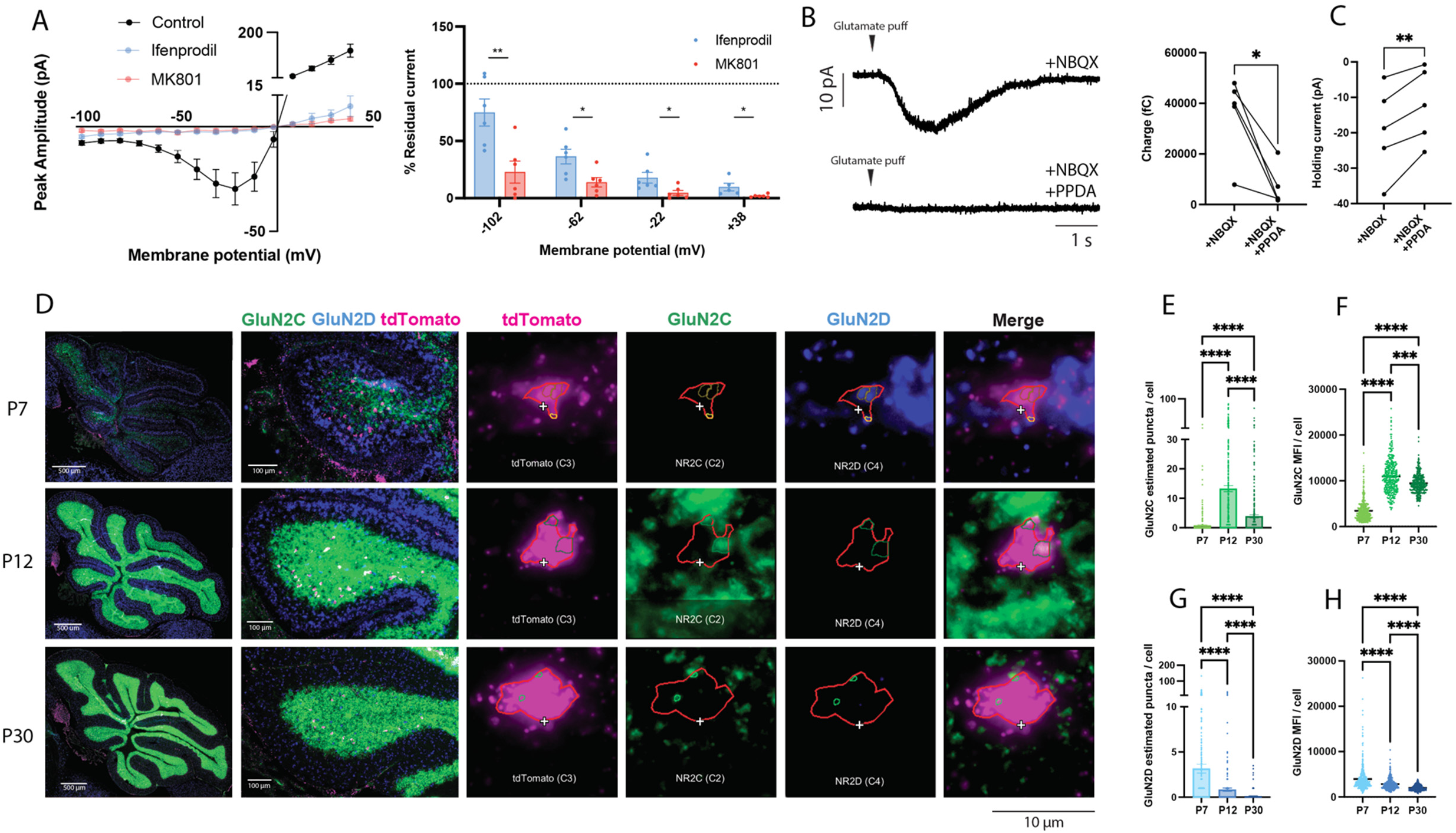
NMDARs in developing UBCs express GluN2C/D subunits A) Left-I-V curves plotting isolated NMDAR-mediated currents evoked by glutamate puffs and the effect of NR2B-selective antagonist (Ifenprodil-blue) or NMDA antagonist (MK801-red). Right-Ifenprodil blocked a larger proportion of the total NMDAR-mediated current at depolarized potentials, consistent with the majority of the current at hyperpolarized potentials being mediated by GluN2C/D subunits. B) GluN2C/D subunit specific antagonist PPDA blocked the NMDAR-mediated current at −75 mV. C) UBCs exhibited GluN2C/D NMDAR-mediated tonic current as seen by outward shift of holding current after application of PPDA. D) RNAscope in-situ hybridization revealed differences in GluN2C and GluN2D expression patterns in P7, P12, and P30 mouse cerebellum. Representative images (from left to right) showing sagittal section of mouse cerebellum, lobe X, and an example each of GluN2C/GluN2D puncta colocalization with tdTomato+ UBCs. E-H) GluN2C expression was higher at P12 and GluN2D expression was higher at P7, measured as either puncta/cell or mean fluorescence intensity (MFI).

To confirm the presence of GluN2C/D subunits in developing UBCs we evoked NMDAR-mediated currents at −75 mV by glutamate puff and measured the pharmacological effect of GluN2C/D subunit selective antagonist, PPDA (1.5 μM) (**Fig** 2B, 79.4 ± 7.93 % inhibition, n = 5; Paired t-test; p = 0.0133) (Feng et al., 2004; Drotos et al., 2025). A tonic NMDAR-mediated current was also revealed by the outward shift in holding current after application of PPDA (**Fig** 2C, 6.93 ± 1.5 pA, n = 5; Paired t-test; p = 0.0097; mean ± SEM).

To corroborate our electrophysiology data, we ran in situ hybridization experiments using RNAScope to visualize and quantify GluN2C/D subunit mRNA in ON UBCs. We used GRP/Ai9 mice which express tdTomato in ON UBCs at three ages – P7, P12, P30. To visualize the change in NMDA mRNA expression across development, we quantified the co-localization of GluN2C and GluN2D puncta with tdTomato mRNA (ON UBCs). We observed GluN2D puncta co-localization with the tdTomato signal at P7 indicating GluN2D expression in P7 ON UBCs. There was considerably less GluN2D expression in ON UBCs at P12 and P30 as reflected in the quantification of estimated puncta/cell (**Fig** 2G) and the mean fluorescence intensity (MFI)/cell (**Fig** 2H). GluN2C expression in ON UBCs was significantly higher at P12 than at P7 or P30, as seen in the quantification of estimated puncta/cell (**Fig** 2E) and the MFI/cell analysis (**Fig** 2F, N = 3-5 animals/condition; n = ∼300 cells/condition; Kruskal-Wallis non-parametric ANOVA test; p < 0.0001).

Taken together, these results show that NMDARs in developing ON UBCs are not only active at depolarized potentials but also at hyperpolarized potentials due to the presence of GluN2C/D subunits. Additionally, these NMDARs mediate a tonic inward current at resting membrane potentials, which although small in amplitude, could play an important role in modulating cell excitability and development because they conduct calcium ions that act as a second messenger for many cellular processes.

### GluN1 KO did not alter UBC brush development

NMDARs are necessary for neuronal development, especially dendritic outgrowth and arborization and synaptic maturation (Kalb, 1994; Rajan and Cline, 1998; Hanson et al., 2019; Hansen et al., 2021). UBCs have a complex dendritic brush that is thought to be critical for their synaptic responses (Rossi et al., 1995; Kinney et al., 1997). We hypothesized that the high expression of NMDARs in developing UBCs play a role in UBC brush development. To test this hypothesis, we established a GluN1 KO model using our GRP/Ai9 mice to prevent the expression of NMDARs in the subset of ON UBCs that were labeled with tdTomato. To validate the KO, we recorded from KO UBCs while stimulating the cell with brief application of 1 mM glutamate (**Fig** 3A) in the presence of AMPA, kainate, mGluR1 and mGluR2/3 antagonists. NMDAR-mediated currents were absent in KO UBCs while control GRP/Ai9 UBCs showed characteristic NMDAR-mediated currents (**Fig** 3B-C).

**Figure 3:**
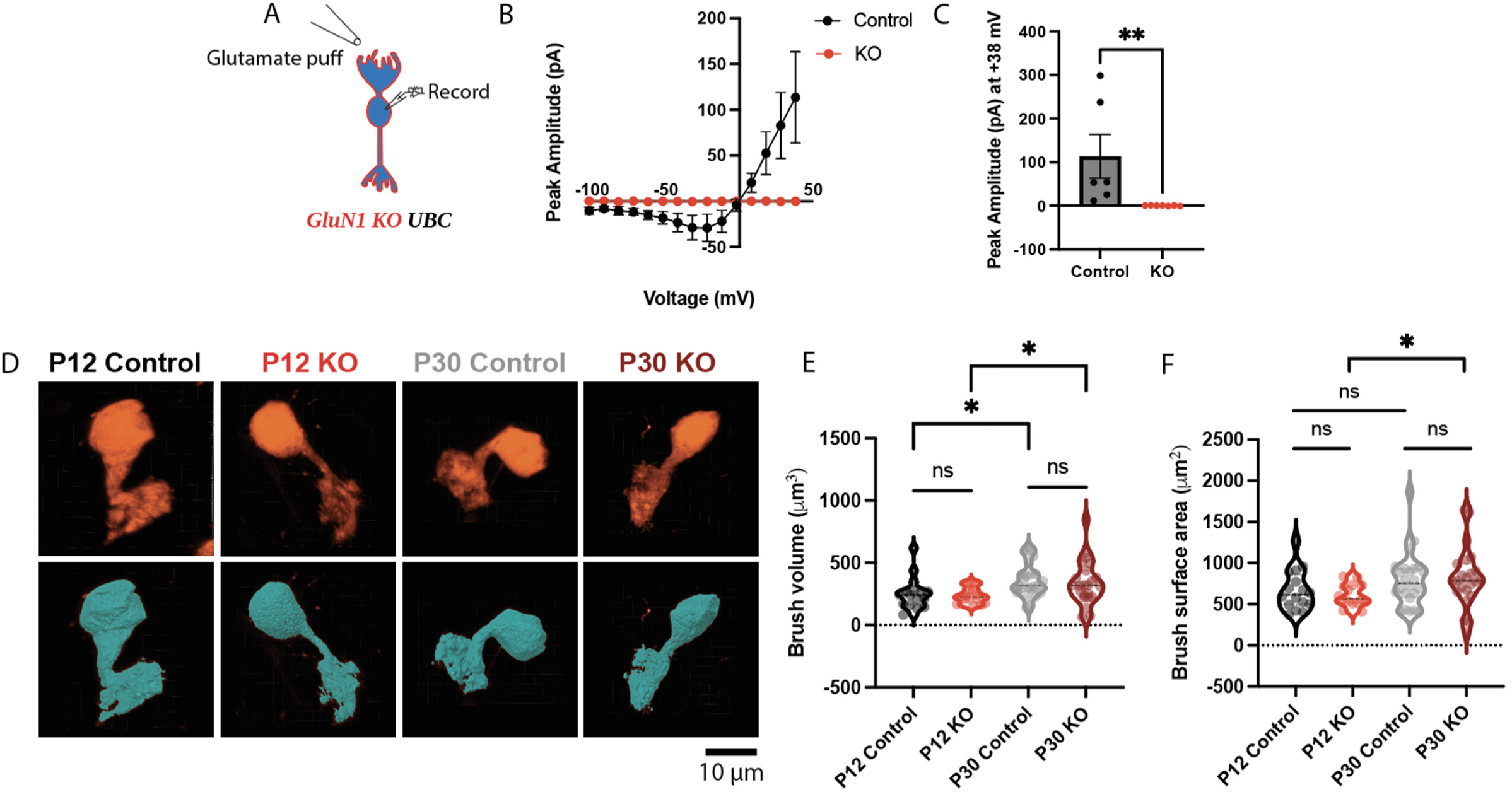
GluN1 KO did not alter UBC brush development A) Illustration of experimental design showing brief (10-30 ms) glutamate puff on GluN1 KO UBC brush while recording from soma. AMPA, kainate, mGluR1 and mGluR2/3 receptors were blocked using 50 μM GYKI53655, 10 μM NBQX, 1 μM JNJ16259685 and 1 μM LY341495, respectively. B) Puffing glutamate while holding the cell at different potentials resulted in a typical NMDA I-V curve in control UBCs while GluN1 KO UBCs showed almost no NMDAR-mediated currents at any potential. C) GluN1 KO UBCs showed successful knock-out of NMDARs as seen by the absence of glutamate puff evoked current at +38 mV. D) Representative confocal images of control and KO UBCs at P12 and P30 (top) and corresponding 3D surface reconstruction (bottom) enabled quantification of brush morphology. E) GluN1 KO did not alter normal brush development or the brush volume (left) and brush surface area (right) at P12 or P30.

Individual control and GluN1 KO UBCs were imaged in slices from P12 and P30 mice using a confocal microscope. 3D surface reconstructions fit to the fluorescence of each cell allowed morphological quantification of the UBC brush (**Fig** 3D). The brush volumes (**Fig** 3E) and surface areas (**Fig** 3F) were not different between control and GluN1 KO UBCs at either age. We did, however, observe an expected increase in brush volume from P12 to P30, in agreement with their incomplete development at P12 and elaboration by late postnatal ages.

### NMDARs regulate UBC number through development

NMDARs have been shown to influence neuronal number during development. Blockade of NMDARs in the developing rat brain triggers widespread apoptosis and decrease in neuronal number (Ikonomidou et al., 1999). In contrast, complete lack of NMDARs results in increased neuronal population in the forebrain of zebrafish (Napoli et al., 2024). Furthermore, cell-death promoting contributions of NMDA signaling cascades are common across neuronal populations (Liu et al., 2007; Martel et al., 2009; Parsons and Raymond, 2014; McQueen et al., 2017).

To explore the role of NMDARs in determining neuronal number, we compared the number of UBCs in lobe X of cerebellum of control and GluN1 KO mice. One side of the cerebellum was sectioned and UBCs were counted from slices 100 µm apart. UBCs were not evenly distributed, as previously reported (Nunzi et al., 2002; Sekerková et al., 2014). There were more UBCs in medial sections than in lateral sections (**Fig** 4A). There was a significant decrease in UBC number from P12 to P30 in control mice (**Fig** 4A, Control P12 vs P30: N = 3-4 mice/condition, 2-way ANOVA; p = 0.0041; **Fig** 4C, top: Welch’s test; p = 0.0490; **Fig** 4C, bottom: Welch’s test; p = 0.0392), but not in GluN1 KO mice (**Fig** 4B, KO P12 vs P30: N=3-4 mice/condition, **Fig** 4B: 2-way ANOVA; p = 0.3419; **Fig** 4C, top: Welch’s test; p = 0.9419; **Fig** 4C, bottom; Welch’s test; p = 0.9417). Note that control and KO mice had the same UBC number at P12 but differ significantly at P30 (**Fig** 4C-D) (Control P12 vs KO P12: (**Fig** 4C, top: Welch’s test; p = 0.6718; **Fig** 4C, bottom: Welch’s test; p = 0.6965); Control P30 vs KO P30: (**Fig** 4C, top: Welch’s test; p = 0.0479; **Fig** 4C, bottom: Welch’s test; p = 0.0136)). We observe the same results when quantifying the density of UBCs in control and KO mice at P12 and P30 (Supplemental **Fig** 1A-C). Area sampled across conditions was not significantly different (Supplemental **Fig** 1D).

**Figure 4:**
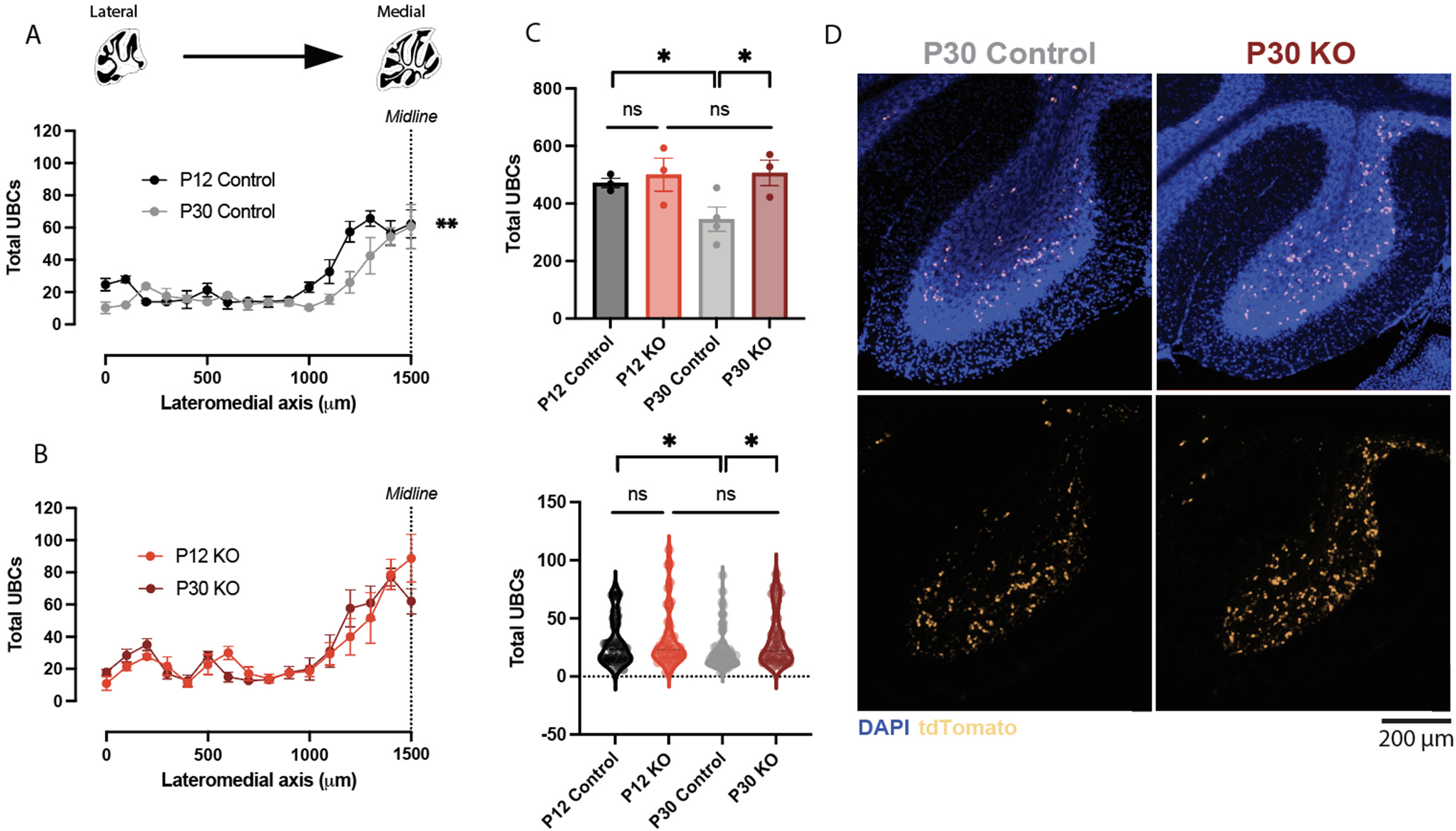
NMDARs regulate UBC number through development A) In control animals there were more UBCs in lobe X at P12 than at P30, indicating a developmental decrease, especially in the medial region. B) Lobe X of GluN1 KO animals had a similar number of UBCs at P12 and P30, suggesting a failure of programmed cell death that occurred in the WTs. C) GluN1 KO did not affect the number of UBCs in lobe X at P12. In control animals, the total number of UBCs in lobe X decreased significantly from P12 to P30. This loss in UBCs was prevented by GluN1 KO. Data are represented per animal (top) and per brain slice (bottom). D) Representative images of lobe X of cerebellum showing fewer UBCs in control animals compared to GluN1 KO animals at P30.

The dorsal cochlear nucleus is a cerebellum-like circuit that also contains a large number of UBCs. GluN1 KO mice had significantly more UBCs in dorsal cochlear nucleus than control mice at P30 (Supplemental **Fig** 1E-F), but not at P12, similar to the cerebellum. Taken together, this provides evidence that NMDARs regulate UBC number in the cerebellum and the dorsal cochlear nucleus, presumably by triggering events that lead to cell death.

### NMDA currents shape evoked firing patterns in ON UBCs

The presence of functional NMDA receptors in developing ON UBCs raises questions about their role in synaptic signaling. It has been previously reported that NMDARs containing GluN2C/D subunits produce particularly slowly decaying EPSPs that contribute to temporal integration of synaptic inputs (Drotos et al., 2025). To test the role of NMDARs in UBC firing, we recorded from developing ON UBCs and stimulated their presynaptic inputs (**Fig** 5A).

**Figure 5:**
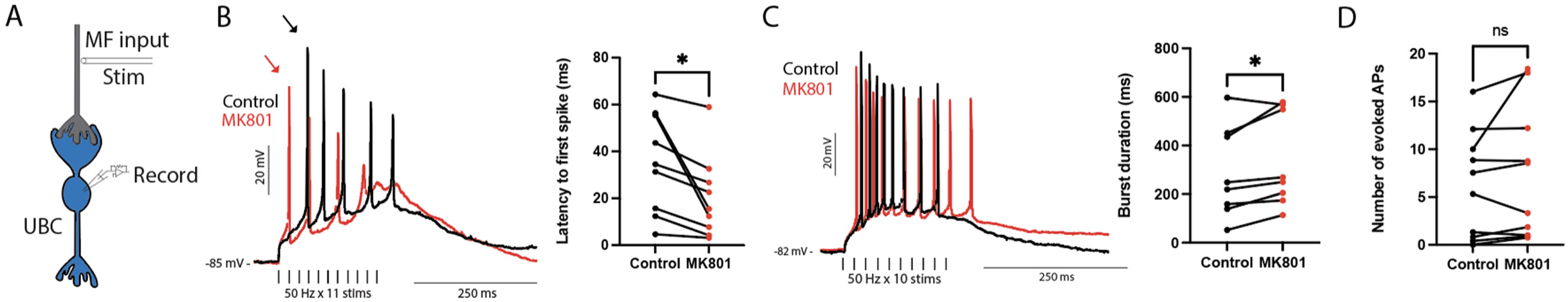
NMDAR-mediated current shapes synaptic stimulation evoked firing in UBCs A) Illustration of experimental design showing MF stimulation while recording from UBC soma. B) Latency to first spike (arrows) evoked by MF stimulation was significantly shorter after blocking NMDARs with MK801 in all UBCs recorded. C) Most cells fired for a longer duration after blocking NMDARs with MK801. D) Blocking NMDARs had no significant effect on the number of spikes evoked.

We predicted that blocking excitatory NMDAR current would increase the latency to the first spike and reduce the firing rate and duration. We were surprised to observe a paradoxical increase in cell excitability after blocking NMDARs. We report two consistent results after blocking NMDARs: a decrease in the latency to the first spike in all cells recorded (**Fig** 5B, Control: 35.37 ± 7.1 ms; MK801: 20.4 ± 5.9 ms; n = 9; Paired t-test; p = 0.0204) and an increase in burst duration (**Fig** 5C, Control: 287.8 ± 66.1 ms; MK801: 338.5 ± 68.6; n = 8; Paired t-test; p = 0.0315). There was no change in the number of action potentials after blocking NMDA receptors (**Fig** 5D, n = 10; Paired t-test; p = 0.2276). Membrane potential was maintained by applying bias current and we confirmed that this prevented it from differing significantly before vs after MK801 application. Input resistance was not affected by MK801 (Supplemental **Fig** 2A-B).

To rule out the possibility of these effects being a result of whole-cell recording for extended periods (intracellular washout or plasticity), we made recordings of synaptic responses for the same duration (over 10 minutes) without application of NMDA antagonist. In these control experiments we did not observe the effects described above (Supplemental **Fig** 3).

Given the presence of a tonic NMDAR-mediated current, we reasoned that these cells could be constantly dampened through this mechanism, which would have a major effect on their intrinsic excitability. We injected the cells with the lowest current that elicited spikes (5 or 10 pA) that we will refer to as rheobase and compared the effect of MK801 on their spiking response. Membrane potential was maintained by applying bias current. A paradoxical increase in excitability after blocking NMDARs was observed, similar to the results using synaptic stimulation. After MK801 application we observed a significant shortening of latency to first spike (**Fig** 6A: Control: 64.29 ± 8.927 ms; MK801: 36.13 ± 4.370 ms; n = 10; Paired t-test; p = 0.0006), significant steepening of the rise slope of depolarization (**Fig** 6B: Control: 0.2741 ± 0.04 mV/ms; MK801: 0.4298 ± 0.05 mV/ms; n = 10; Paired t-test; p = 0.0004), and a significant increase in firing rate on rheobase current injection (**Fig** 6C: Control: 20.02 ± 5.098 Hz; MK801: 31.04 ± 6.745 Hz; n = 9; Paired t-test; p = 0.0068). Input resistance did not change significantly on MK801 application (Supplemental **Fig** 2C-D).

**Figure 6:**
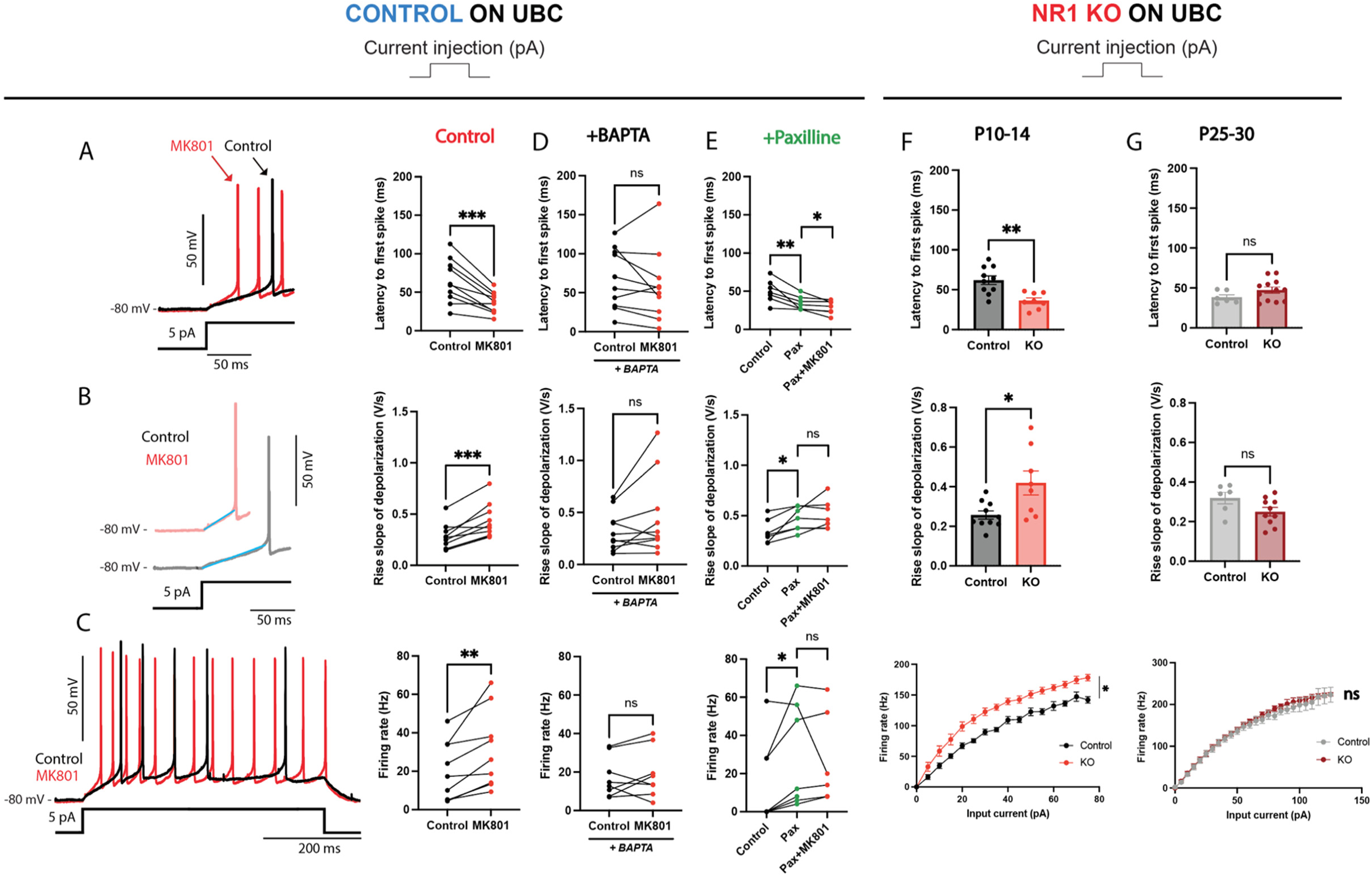
Dampening effect of NMDARs is calcium dependent and is occluded by BK channel blockade A) Decrease in latency to first spike (arrows) after blocking NMDARs occurs with current injections, consistent with the block of a tonic NMDA current. B) The rise slope of depolarization on current injections became steeper after blocking NMDARs. C) Blocking NMDARs results in increased excitability as seen by increase in firing rate. D) Effects of blocking NMDARs on latency, rise slope of depolarization and firing rate (top, middle and bottom, respectively) are prevented by presence of BAPTA in the internal pipette solution. E) Presence of Paxilline in the bath occluded the effect of NMDAR block partially on latency (top) and completely on rise slope of depolarization (middle) and firing rate (bottom) suggesting BK channels might be downstream of NMDA activity. F) GluN1 KO had a dampening effect that was similar to pharmacological blockade. Following current injections, latency to first spike was significantly shorter (top), rise slope of depolarization was significantly greater (middle) and firing rate was significantly higher (bottom) in GluN1 KO UBCs compared to control UBCs. G) Effects seen in (F) in developing animals were not present at P25-30 consistent with a decrease in NMDAR-mediated currents at this age.

### Dampening effect of NMDA receptors is calcium dependent and is occluded by BK channel inhibition

NMDARs are permeable to calcium, so we tested whether the dampening effect after NMDAR blockade was calcium mediated. We repeated the rheobase current injection experiments (**Fig** 6A-C) with high-affinity calcium chelator BAPTA (15 mM) in the pipette. Blocking NMDARs with MK801 with BAPTA in the pipette prevented the change in the latency to first spike (**Fig** 6D, top; n = 10, Paired t-test; p = 0.2860), rise slope of depolarization (**Fig** 6D, middle; n = 10, Paired t-test; p = 0.0989) and firing rate (**Fig** 6D, bottom; n = 10, Paired t-test; p = 0.4743). These results suggest that calcium flux through NMDARs is responsible for the effect on excitability.

NMDAR-mediated calcium influx can activate calcium activated potassium channels in neurons (Nicoll and Alger, 1981; Isaacson and Murphy, 2001). Functional coupling of tonically active NMDARs with calcium activated potassium channels has been hypothesized to result in dampening of cell excitability in a computational model (Gall and Dupont, 2019). Large conductance calcium and voltage activated potassium (BK) channels are one such type of calcium activated potassium channels which can modulate firing rates in neurons (Marty, 1981; Gu et al., 2007). UBCs have been reported to have large BK channel currents, which are coupled with NMDARs in various neuronal cell types (Isaacson and Murphy, 2001; Diana et al., 2007; Zhang et al., 2018; Martínez-Lazaro et al., 2025).

To test if BK channels are necessary for NMDAR-mediated dampening of cell excitability, we repeated the current injection experiments (**Fig** 6A-C) in the presence of 10 μM Paxilline, a selective BK channel blocker. We predicted that Paxilline would occlude the effect of MK801 on latency to first spike, rise slope of depolarization and firing rate. Paxilline alone decreased the latency to first spike (**Fig** 6E, top, n = 7; Paired t-test; p = 0.0073), steepened the rise slope of depolarization (**Fig** 6E, middle, n = 7; Paired t-test; p = 0.0102) and increased firing rate (**Fig** 6E, bottom, n = 7; Paired t-test; p = 0.0494). This result supports the presence of a tonic calcium current in UBCs that activates BK channels. Blocking NMDARs with MK801 had little effect in the presence of Paxilline. Latency to first spike decreased further (**Fig** 6E, top, n = 7; Paired t-test; p = 0.04), but the rise slope of depolarization (**Fig** 6E, middle, n = 7; Paired t-test; p = 0.2670) and firing rate (**Fig** 6E, bottom, n = 7; Paired t-test; p = 0.5282) remained unaffected. Changes in latency to first spike, rise slope of depolarization and firing rate were similar with MK801 application alone, Paxilline application alone and MK801 application in the presence of Paxilline (Supplemental **Fig** 4). These results suggest that BK channels mediate the effects of NMDAR blockade on UBC firing.

If NMDARs dampen excitability, then the GluN1 KO UBCs may be more excitable. At P10-14, compared to controls, KO UBCs showed a shorter latency to first spike (**Fig** 6F, top; Control: 61.96 ± 5.432 ms, n = 10; KO: 36.21 ± 3.578 ms, n = 8; Welch’s test; p = 0.0013), steeper rise slope of depolarization (**Fig** 6F, middle; Control: 0.2569 ± 0.021 mV/ms, n = 10; KO: 0.4188 ± 0.061 mV/ms, n = 8; Welch’s test; p = 0.0332) and higher firing rate at almost all input currents tested (**Fig** 6F, bottom; Control: n = 16; KO: n = 10; Mixed-effects ANOVA; p < 0.0001), remarkably similar to the effect of NMDA receptor blockade in wild types. In contrast, at P25-30, there were no significant differences between control and KO UBCs with regard to latency to first spike (**Fig** 6G, top; Control: 38.20 ± 3.204 ms, n = 6; KO: 47.29 ± 3.92 ms, n = 12; Welch’s test; p = 0.0921), rise slope of depolarization (**Fig** 6G, middle; Control: 0.3191 ± 0.03 mV/ms, n = 6; KO: 0.2498 ± 0.022 mV/ms, n = 10; Welch’s test; p = 0.0876) or firing rate (**Fig** 6G, bottom; Control: n = 11; KO: n = 14; Mixed-effects ANOVA; p = 0.9496), consistent with a lack of NMDAR-mediated currents at those ages in WT UBCs. In summary, our data suggest that NMDARs in developing UBCs dampen excitability in a calcium dependent manner, likely via calcium activated potassium channels.

### Effects of tonic NMDA currents on UBC firing can be replicated in a computational model

To explore the conditions under which tonic NMDA and BK conductances can produce the paradoxical dampening effect of NMDARs that we observed, we built a single compartment model of a UBC using NEURON (Hines and Carnevale, 1997; Carnevale, 2005). We injected a small current into the UBC model and measured the latency to first spike (18.2 ms), the rise slope of depolarization (1.09 mV/ms) and total spikes evoked (5 spikes) (**Fig** 7A). We then repeated this simulation with the conductance of NMDARs reduced to zero to reflect NMDAR block with MK801. As in our experimental data, we saw a significant shortening of latency to first spike (8.8 ms), steepening of the rise slope of depolarization (2.63 mV/ms) and an increase in the total number of spikes (10 spikes).

**Figure 7:**
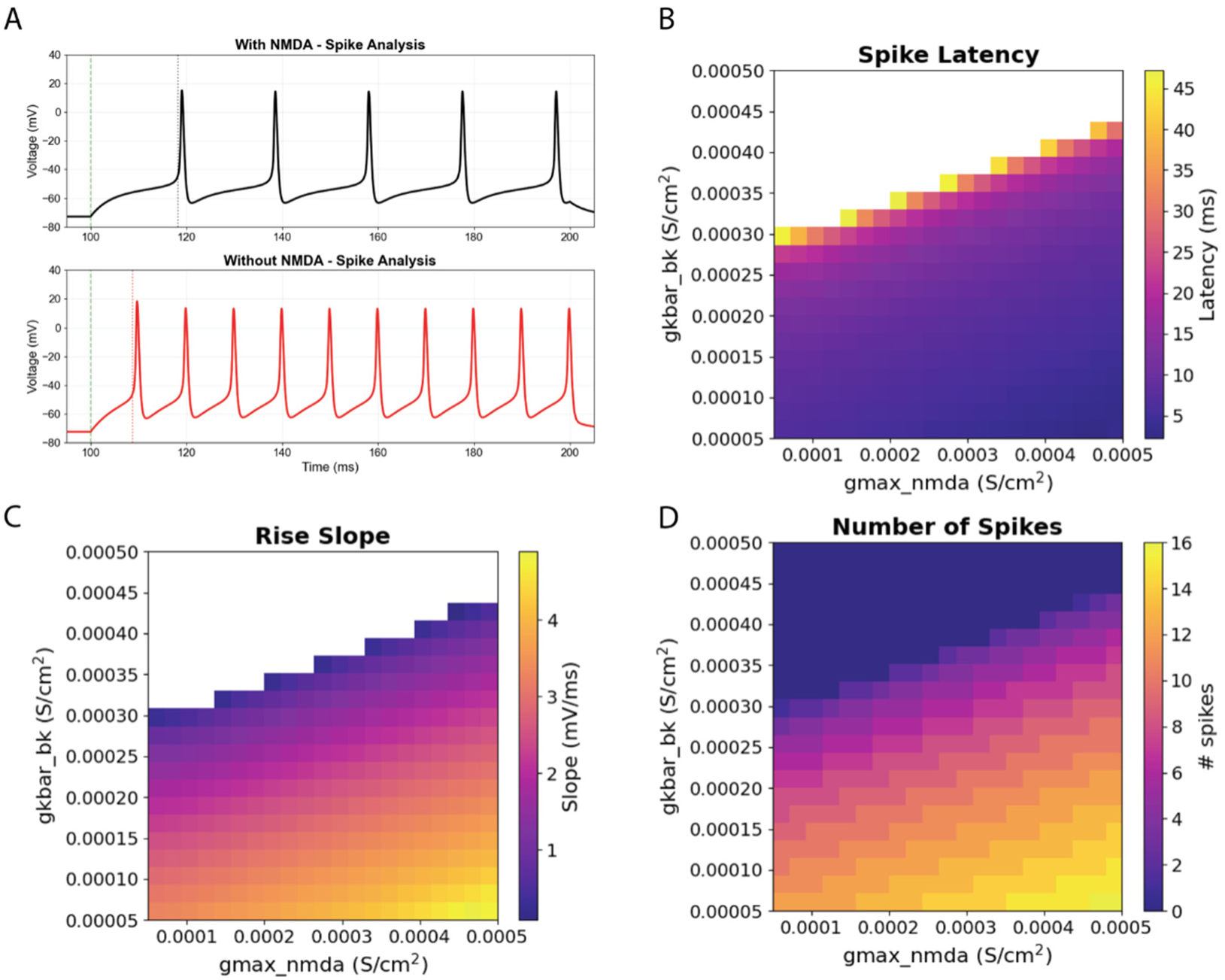
Effects of NMDAR-mediated currents on UBC firing can be independently replicated using a computational model of UBC A) Blocking NMDARs in a computational model replicated the effects observed experimentally on latency to first spike, rise slope of depolarization, and firing rate (Figure 5E, F and G). Latency decreased from 18.2 ms to 8.8 ms. Rise slope of depolarization increased from 1.09 mV/ms to 2.63 mV/ms. Number of spikes increased from 5 spikes to 10 spikes. Dotted green line represents the start of stimulus. Dotted black or red line represent onset of first spike (when dV/dt crossed 10 mV/ms). B-D) 2D parameter sweeps plotted as heatmaps revealed the range over which NMDA and BK conductances could affect latency to first spike (B), the rise slope of depolarization (C), and the number of spikes (D). White spaces indicate the cell did not fire with those conductance pairs.

Parameter sweeps from conductances 0.00005 to 0.0005 S/cm^2^ for NMDA and BK were used to test the range over which the spike latency, rise slope, and spike rate varied, and the results were plotted as a heatmap (**Fig** 7B-D). White spaces indicate the cell did not fire with those conductance pairs. Spike latency increased with both NMDA and BK conductances (**Fig** 7B). The rise slope of depolarization was steepest with high NMDAR and low BK conductance (**Fig** 7C). The fastest firing was elicited when BK conductance was low and NMDA conductance was high (**Fig** 7D). The model was robust over a wide range of conductances and supports the hypothesis that tonic NMDA receptor activity dampens excitability by increasing a calcium activated potassium current.

## DISCUSSION

In this study, we report that developing (P10-14) UBCs in mouse cerebellum have significant expression of NMDARs which decrease by P25-30. We show that developing ON UBCs have GluN2B and GluN2C/D subunits which generate a tonic NMDA current at resting membrane potentials. NMDARs regulate the number of ON UBCs in P25-30 mouse cerebellum. Surprisingly, NMDARs dampen the excitability of developing ON UBCs in a calcium dependent manner, likely mediated by calcium activated potassium channels.

### Developmental expression of NMDARs in UBCs

NMDAR contributions to synaptic signaling have been suggested to decline with age in a study using UBCs from P10-30 rats, but a quantitative comparison across ages has not been reported (Rossi et al., 1995). A recent study showed that NMDAR had no significant contributions to UBC firing evoked by synaptic stimulation in P30-47 mice (Huson and Regehr, 2024). In contrast, NMDAR-mediated currents have been shown in UBCs from P8-20 rats and P20-80 mice (Billups et al., 2002; Canton-Josh et al., 2022). In the present study we found no evidence of NMDARs in P25-30 ON UBCs, while P25-30 OFF UBCs had some NMDAR-mediated currents which were smaller than at P10-14. Consistent with downregulation of NMDARs in mature UBCs, we saw significant differences in UBC excitability in GluN1 KO UBCs at P10-14 compared to age-matched controls, but no differences between KO and controls at P25-30. Thus, NMDARs are present during development and have a role in synaptic signaling but are lost during development in the ON UBCs studied.

We used ON UBCs labeled in one transgenic mouse line (GRP-Cre), which is a subset of the ON UBCs in the mouse cerebellum and dorsal cochlear nucleus (Hariani et al., 2024). We used this mouse line because it allowed genetic deletion of NMDARs in ON UBCs specifically, although this limited our ability to test other UBC types. Recent studies have suggested a continuum of UBC types based on their gene expression and synaptic responses recorded extracellularly (Guo et al., 2021; Kozareva et al., 2021; Huson et al., 2023). In this study we showed that UBCs of all types have significant NMDAR-mediated currents at early postnatal ages (P10-14) and, although GRP ON UBCs lose them entirely, other types may maintain them into maturity.

### Effect of GluN1 KO in ON UBC dendritic brush development

GluN1 are obligatory components of NMDAR assembly and thus GluN1 KO results in complete lack of NMDARs (Forrest et al., 1994). The GluN1 KO UBCs had similar electrophysiological features compared to control UBCs at P10-14 and P25-30, including cell capacitance, input resistance, and voltage sag. We observed similar significant changes in intrinsic properties from P10-14 to P25-30 in control and KO UBCs (increase in capacitance, decrease in input resistance, and increase in voltage sag) (Supplemental **Fig** 5). In several neuron types, NMDARs promote dendritic outgrowth and arborization (Kalb, 1994; Rajan and Cline, 1998; Lee et al., 2005). GluN2C/D-NMDAR-mediated currents in particular have been shown to facilitate dendritic and synaptic maturation in cortical interneurons (Hanson et al., 2019). Therefore, it was surprising that GluN1 KO UBCs developed dendritic brushes that could not be distinguished from those of control UBCs. It is possible that there were differences in fine structure that could not be resolved with confocal microscopy. Alternatively, UBCs may compensate for GluN1 KO by expression of other ion channels to promote normal brush development. Such compensation of ion channels occurs in other neuron types to maintain cell excitability (Pratt and Aizenman, 2007; Kulik et al., 2019; Karmelic et al., 2026). NMDARs are thought to guide dendritic development through their signaling that leads to an influx of calcium (Konur and Ghosh, 2005; Lau et al., 2009; Rosenberg and Spitzer, 2011). Developing UBCs may have other significant non-NMDA sources of calcium influx that govern dendritic development, such as voltage gated calcium channels, or calcium permeable AMPA receptors, the presence of which has not been tested in developing UBCs (Burnashev et al., 1992; Diana et al., 2007). Additional experiments are necessary to determine what guides the development of the UBC’s unusual dendritic brush.

### NMDARs regulate ON UBC number

We found higher UBC numbers in lobe X and dorsal cochlear nucleus of GluN1 KO mice compared to control mice at P30. Two hypotheses could explain our data. First, more UBCs are born and migrate to the cerebellum, or second, fewer UBCs undergo cell death through development. GluN1 KO has been shown to delay maturation of transit-amplifying neuroblasts into post-mitotic neurons, which results in increased neuronal populations in forebrain of zebrafish (Napoli et al., 2024). However, we see no differences in UBC numbers between control and KO mice earlier, at P12. This suggests birth and migration of similar numbers of UBCs into lobe X of cerebellum and a possible role of cell autonomous NMDAR activity through development to control UBC numbers. Activating NMDARs with GluN2B subunits has been shown to result in excitotoxicity and increased apoptosis (Liu et al., 2007; Parsons and Raymond, 2014). Lack of such cell death cascades in GluN1 KO UBCs during development could result in increased UBC numbers in adulthood. Future studies that quantify apoptosis will be required to test this hypothesis.

### Effect of NMDAR activity on UBC excitability

NMDARs typically have a direct depolarizing effect that increases excitability. In contrast, we observed a dampening of excitability by NMDARs in experiments in which depolarization was driven by current injection or by synaptic stimulation. The result that block of NMDAR increased excitability in response to direct depolarization by current injection is consistent with a NMDAR-mediated tonic current. We previously showed that ambient glutamate is present at the mossy fiber-UBC synapse and it generates tonic AMPA and mGluR2 currents (Balmer et al., 2021). GluN2C/D containing NMDARs are likely to produce a tonic inward current in the presence of ambient glutamate as well, because of their relative insensitivity to magnesium block, slow and incomplete desensitization, and high glutamate affinity (Hollmann and Heinemann, 1994; Vicini et al., 1998; Qian et al., 2005; Meur et al., 2007; Paoletti et al., 2013; Hanson et al., 2019; Mao et al., 2020).

Constitutive deletion of NMDARs in the GluN1 KO mouse produced effects that were recapitulated by pharmacological blockade of NMDARs in control mice. Latency to first spike was shortened, rise slope of depolarization was increased, and the frequency-intensity curves of the GluN1 KO UBCs showed higher overall excitability compared to control developing UBCs. This suggests that the effect was due to a lack of NMDA receptors rather than to altered development of the cells. Similar hyperexcitability after NR1 KO or pharmacological inhibition of NMDAR has been observed in dorsal root ganglion neurons, which was suggested to be due to activation of SK channels (Pagadala et al., 2013). It is surprising that there was no apparent compensation by other channels to maintain the excitability of these cells, which occurs in many other cell types (Pratt and Aizenman, 2007; Kulik et al., 2019; Karmelic et al., 2026). No differences were observed in these measures in older GluN1 KO UBCs (P25-30), consistent with a lack of NMDARs at this age in controls.

In occlusion experiments with Paxilline, MK801 had a significant, but greatly reduced, effect on latency to first spike even in the presence of Paxilline. This suggests that BK channels might not be the only mediator of NMDAR-mediated inhibition. Small potassium (SK) channels have been shown to be coupled with NMDARs in other neurons (Faber, 2010; Pagadala et al., 2013; Ferreira-Neto and Stern, 2021). Many other channels can be modulated by calcium influx, for example, Kv4 channels mediating the ‘A’ current (An et al., 2000). These potassium channels may be additional downstream effectors of NMDARs in developing UBCs. However, our data provides evidence that BK channels mediate the majority of the effect of NMDAR blockade on excitability.

### Potential effects of tonic NMDA-BK activity

NMDA receptors are coupled to calcium-activated potassium channels in numerous brain areas including hippocampus, olfactory bulb, cerebellum, cortex and thalamus (Zorumski et al., 1989; Isaacson and Murphy, 2001; Zhang et al., 2018; Gómez et al., 2021). In layer 5 neurons in the barrel cortex, NMDAR-BK coupling raises the threshold for synaptic plasticity and confers pathway-specific plasticity (Gómez et al., 2021). Our findings suggest that a similar mechanism may operate in UBCs, where NMDA mediated dampening of excitability may regulate synaptic plasticity. Although tonic NMDA currents have been observed in several neuronal types (Sah et al., 1989; Meur et al., 2007; Povysheva and Johnson, 2012; Hanson et al., 2019; Kim et al., 2024), to our knowledge, tonic activation of coupled calcium activated potassium channels has not been reported. Computational studies predict that tonic NMDA-BK activity promotes bistability in neuronal excitability by dramatically extending the range of stimulation where quiescence and stable firing can coexist (Gall and Dupont, 2019). Such bistability has long been proposed to support the encoding of transient synaptic events and contribute to memory formation (Goldbeter, 1996; Marder et al., 1996). In UBCs, tonic NMDA-BK signaling may similarly broaden the computational processing ability thereby influencing the transformation of vestibular signals that underlie balance and eye movements.

## Data Availability

The data that support the findings of this study are available upon reasonable request.

---

**Supplemental Figure 1:**
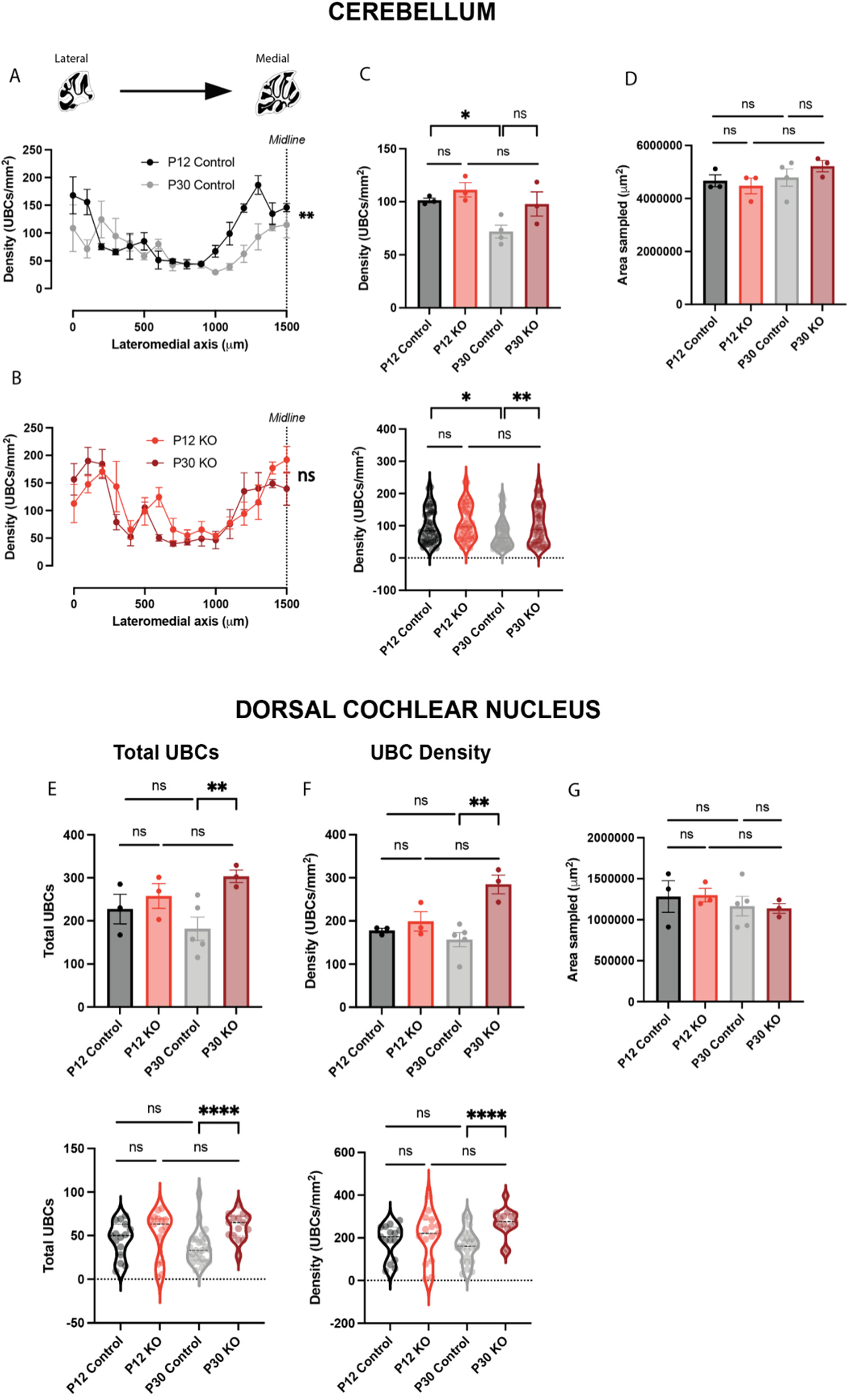
NMDARs regulate total number and density of UBCs in the cerebellum and the DCN A) Lobe X of control P12 animals had greater density of UBCs as compared to control P30 animals. B) Lobe X of GluN1 KO animals show similar density of UBCs at P12 and P30. C) GluN1 KO does not affect the density of UBCs in lobe X at P12. In control animals, density of UBCs in lobe X decreases significantly through development to P30. This loss in density of UBCs is prevented by GluN1 KO. Data represented per animal (top) and per slice (bottom). D) The area of lobe X sampled per condition was not significantly different. E-F) Increase in total UBC number (E) and density (F) is observed in the DCN at P30 in GluN1 KO animals, as observed in the cerebellum. G) The area of DCN sampled per condition was not significantly different.

**Supplemental Figure 2:**
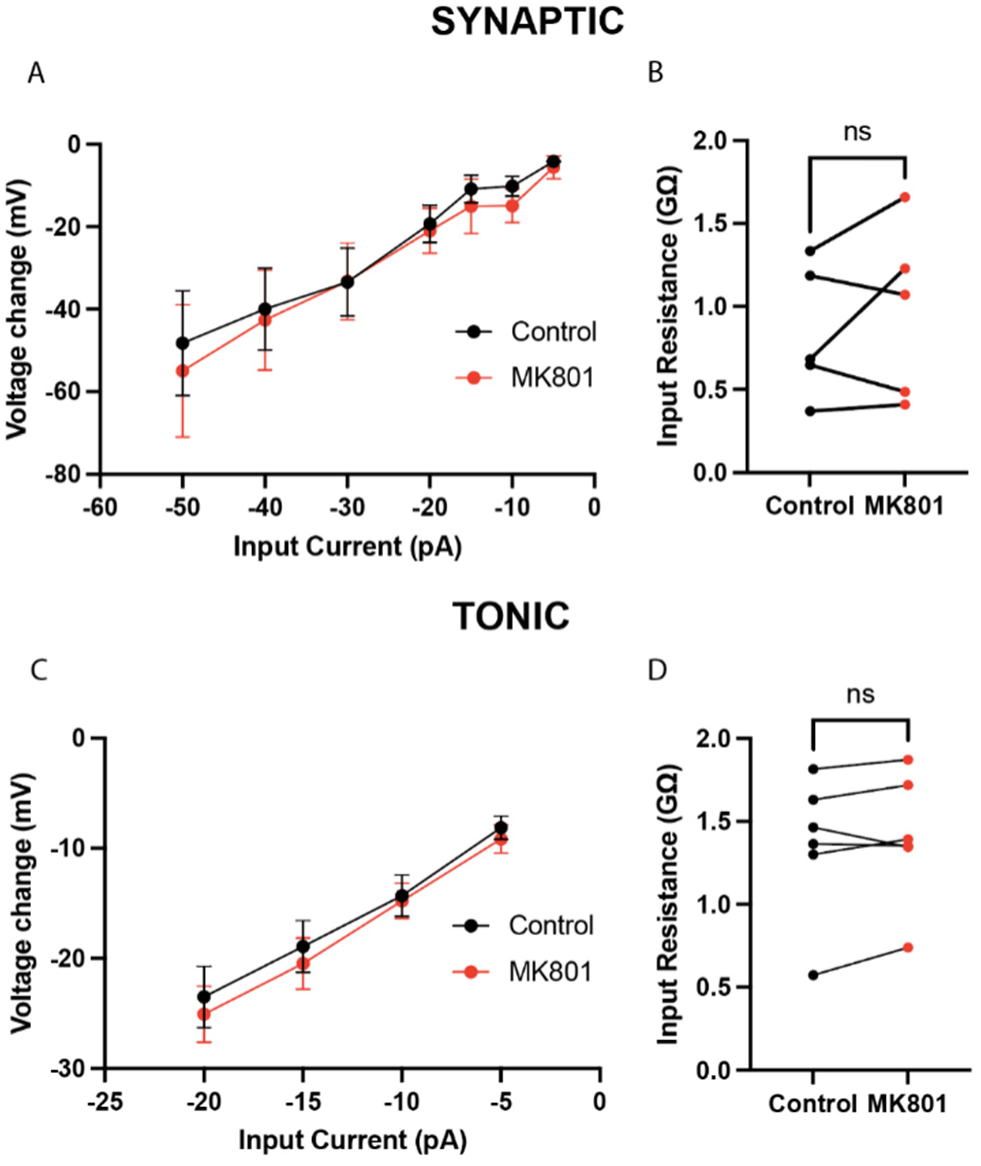
Input resistance was not affected by blocking NMDARs with MK801. A) Plotting the I-V curve for hyperpolarizing current steps revealed no change in input resistance after blocking NMDARs with MK801. These cells are the same as used for Figure 5B-D. B) Input resistance calculated by Clampfit software also showed no change after blocking NMDARs with MK801. C) Similar to (A), no change in input resistance was observed in the I-V curve plotted for cells used in Figure 5E-G. D) Input resistance calculated by Clampfit software also showed no change after blocking NMDARs with MK801.

**Supplemental Figure 3:**
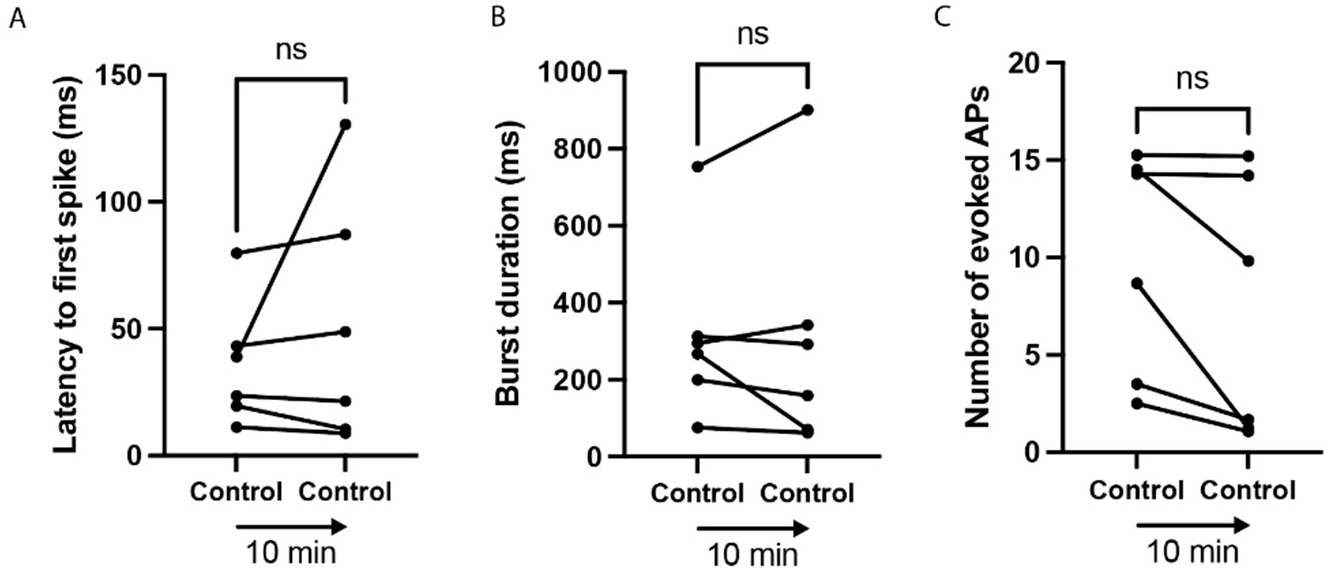
Control experiment to rule out whole-cell recording as cause of altered UBC firing pattern A-C) Recording the cell for 10 minutes did not change the latency to first spike (A), the burst duration (B) or the number of spikes evoked (C) after synaptic stimulation of presynaptic MF.

**Supplemental Figure 4:**
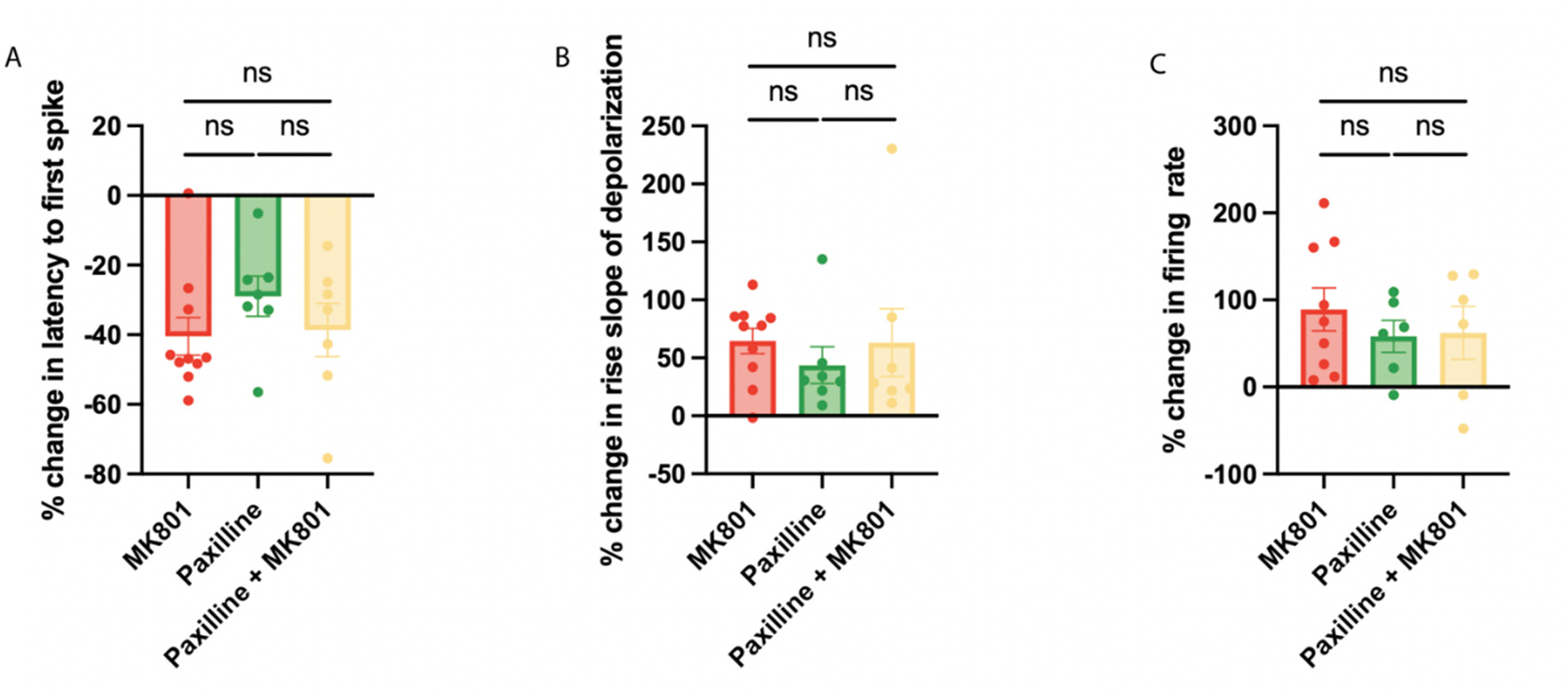
Paxilline occludes the effect on MK801 on UBC firing patterns A-C) Paxilline occluded the effect of MK801 on UBC firing patterns. MK801 did not change the latency to first spike (A), rise slope of depolarization (B) or the firing rate (C) significantly in the presence of Paxilline.

**Supplemental Figure 5:**
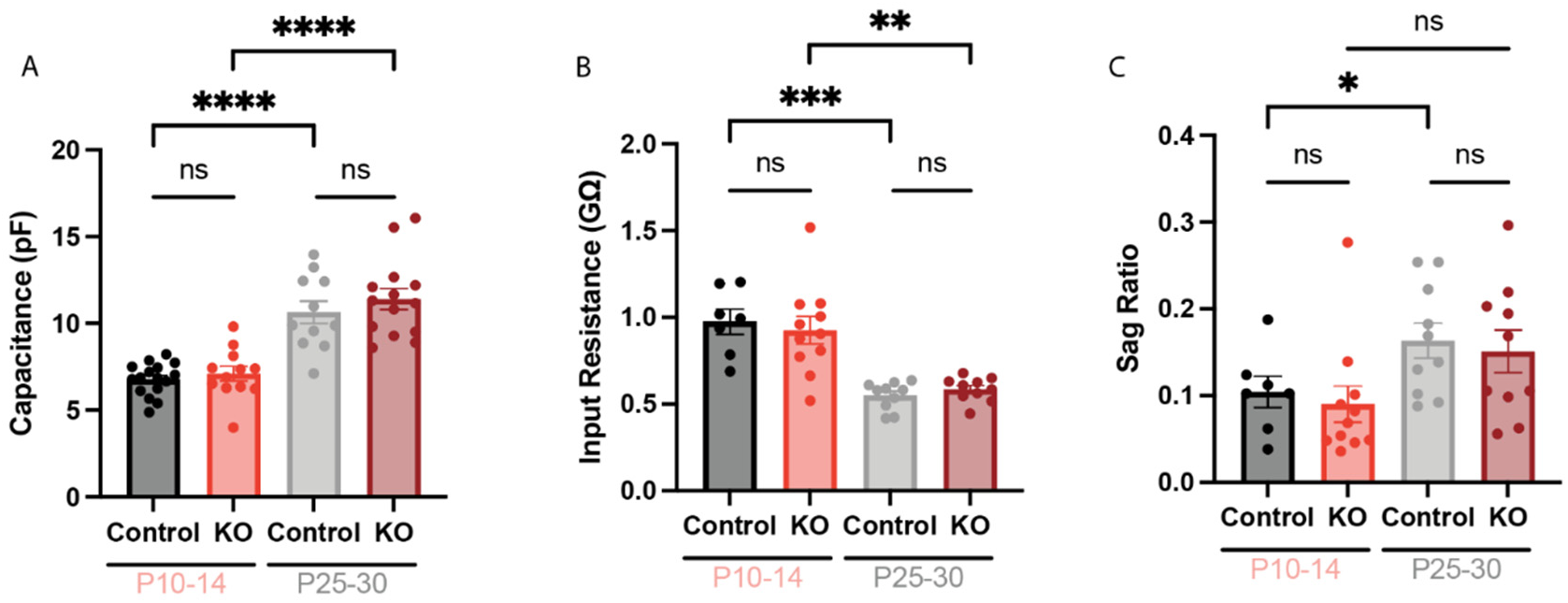
Intrinsic properties of WT and GluN1 KO UBCs at different ages (A) Capacitance was similar for control and GluN1 KO UBCs. However, it increased through development. (B) Input resistance did not change with GluN1 KO but decreased through development. (C) Sag Ratio remained comparable after GluN1 KO but increased through development.

